# Development Of Novel Multiplex PCR For Rapid Diagnosis Of Coinfected Hemo-Parasites In Cattle

**DOI:** 10.1101/2021.06.14.448329

**Authors:** Pankaj Kumar, Abhay Kumar, Kamal Sarma, Paresh Sharma, Rashmi Rekha Kumari, Manish Kumar

## Abstract

A novel, rapid and specific multiplex polymerase chain reaction has been developed for the diagnosis of hemo-parasitic infection in bovine blood by three of the most common hemo-parasites. The reported method relied on the detection of the three different bovine hemoparasites isolated from red blood cells (RBCs) of cattle by conventional Giemsa stained blood smear (GSBS) and confirmed by multiplex PCR. The designed multiplex primer sets can amplify 205, 313 and 422 bp fragments of *apocytochrome b, sporozoite and macroschizont 2 (spm2)* and *16S rRNA* gene for *Babesia bigemina, Theileria annulata* and *Anaplasma marginale*, respectively. This multiplex PCR was sensitive with the ability to detect the presence of 150 ng of genomic DNA. The primers used in this multiplex PCR also showed highly specific amplification of specific gene fragments of each respective parasite DNA without the presence of non-specific and non-target PCR products. This multiplex PCR system was used to diagnose GSBS confirmed blood samples (N=12) found infected or co-infected with hemoparasites. A comparison of the two detection methods revealed that 58.33% of specimens showed concordant diagnoses with both techniques. The specificity, positive predictive value and kappa coefficient of agreement was highest for diagnosis of *B. bigemina* and lowest for *A. marginale*. The overall Kappa coefficient for diagnosis based on GSBS for multiple pathogen compared to multiplex PCR was 0.56 slightly behind the threshold of 0.6 of agreement. Therefore, confirmation should always be made based on PCR to rule out false positive due to differences in subjective observations, stain particles and false negative due to low level of parasitaemia. The simplicity and rapidity of this specific multiplex PCR method make it suitable for large-scale epidemiological studies and for follow-up of drug treatments.

## Introduction

Recent studies suggest that vector borne diseases has emerged due to climate change in tropical countries (Balakrishnan 2017; El-Sayed and Kamel 2020; Kumar et al. 2021; Kumari et al. 2019). Tick population has increased considerably and so is the tick borne infections in susceptible host. It was observed that cattle population often suffer from coinfection of more than one pathogens transmitted by the ticks. These infections cannot be diagnosed symptomatically as the symptoms are overlapping and elusive. Similarly microscopic examination based on conventional blood smear examination often requires expertise observation and may sometimes be misleading in co-infections or due to subjective observations. PCR based diagnosis is very specific and sensitive in detecting these pathogens, however may require multiple PCR for diagnosis of pathogen load in samples suspected to have co-infections. Considering these limitations of diagnosis of multiple tick transmitted diseases, an attempt has been made to design a multiplex PCR to detect co-infections of these pathogens in a single run with the objective to provide prompt and complete diagnosis to the farmers.

## Materials and Methods

### Ethics

The study involved drawing of 3 ml blood from jugular vein aseptically from clinically affected animal with the consent of the owners. These were either brought for treatment to veterinary hospitals or as a call for treatment at farmer’s door by the local veterinarian. There are no specific ethical guidelines for a blood sample collection from clinical cases, and hence no prior approval was mandatory. Treatment was provided to all animals sampled.

### Animal

Blood samples were selectively collected from crossbred cattle (*Bos taurus* × *Bos indicus* breeds) (n=30) in an EDTA vacutainer tube and a clot activator tube (BD, Franklin, USA) suspected to have infection of tick transmitted diseases. Cattle were chosen from the farmer’s field, dairy farms and veterinary hospitals around endemic pockets of Manner block of the peri-urban Patna located in the Gangetic plains of Bihar, India. The selection of cattle was made on the available clinical history and symptoms associated with tick infestation.

### Clinical examination

Selected animals from the endemic area were clinically examined during sampling. Rectal temperature for understanding the severity, mucosal membranes for status of anemia and pre-scapular lymph nodes for any enlargement, respiration pattern was also observed. Eye was examined for any opacity or abnormal discharge. Any other abnormality observed during clinical examination was also recorded.

### Blood examination

The collected blood samples were processed for microscopic examination, hematological examination, and whole blood genomic DNA isolation. For microscopic examination of blood and lymph-node aspirate, a duplicate thin smear was prepared on a glass slide from each sample and fixed with methanol for 5 min. The fixed smear on the glass slide was stained with Giemsa stain (Himedia) for 20 min, washed, air-dried, and examined under oil (100x) immersion. The presence of piroplasms in blood smear through microscopic examination was considered positive for *Theileria* spp. The hemoglobin (Hb) concentration (g/dL) in blood was measured using HemoCue^®^ cuvettes and its analyzer. PCV was calculated based on correlation between hemoglobin and PCV expressed as Hb concentration (g/dL) = 0.3 PCV + 3 (Turkson and Ganyo 2015). RBC count and WBC counts were measured manually under microscope by routine procedures (Jain 1999).

### DNA Isolation

Infected blood samples (300 μL) found positive based on GSBS were used to extract whole genomic DNA using a commercial kit (GCC) as per the manufacturer’s instruction and stored at −20°C. The quality and quantity of genomic DNA were checked by running the extracted DNA on agarose gel (1%), and the purity was measured using nanodrop readings.

### Primer Designing

The primer was designed for identification of three most common pathogens (*Theileria annulata, Anaplasma marginale* and *Babesia bigemina*) in bovine based on our experience and infection prevailing in the region (Table 1). The primers of all the three pathogens was designed to get nearly similar melting temperature (Tm~57-59°C). Primer for *T. annulata* (313 bp) was designed using NCBI reference sequence XM_947453.1 for 2763 bp sequence of *T. annulata* Spm2 protein partial mRNA (Pain et al., 2005). The *sporozoite and macroschizont 2* (*spm2*) gene sequence is reported to be specific to *T. annulata* parasitic infection (Prabhakaran et al. 2021; Tian et al. 2018). Primer for simultaneous detection of *A. marginale* (422 bp) was designed using NCBI reference sequence for 16S ribosomal RNA sequence of *A. marginale* str. Florida as Gene ID-7398331. The primer for the third common pathogen *B. bigemina* (205 bp) was designed using NCBI reference nucleotide sequence AF109354.1 for 720 bp of *apocytochrome b* gene (Bilgiç et al. 2013).

**Table 1.**
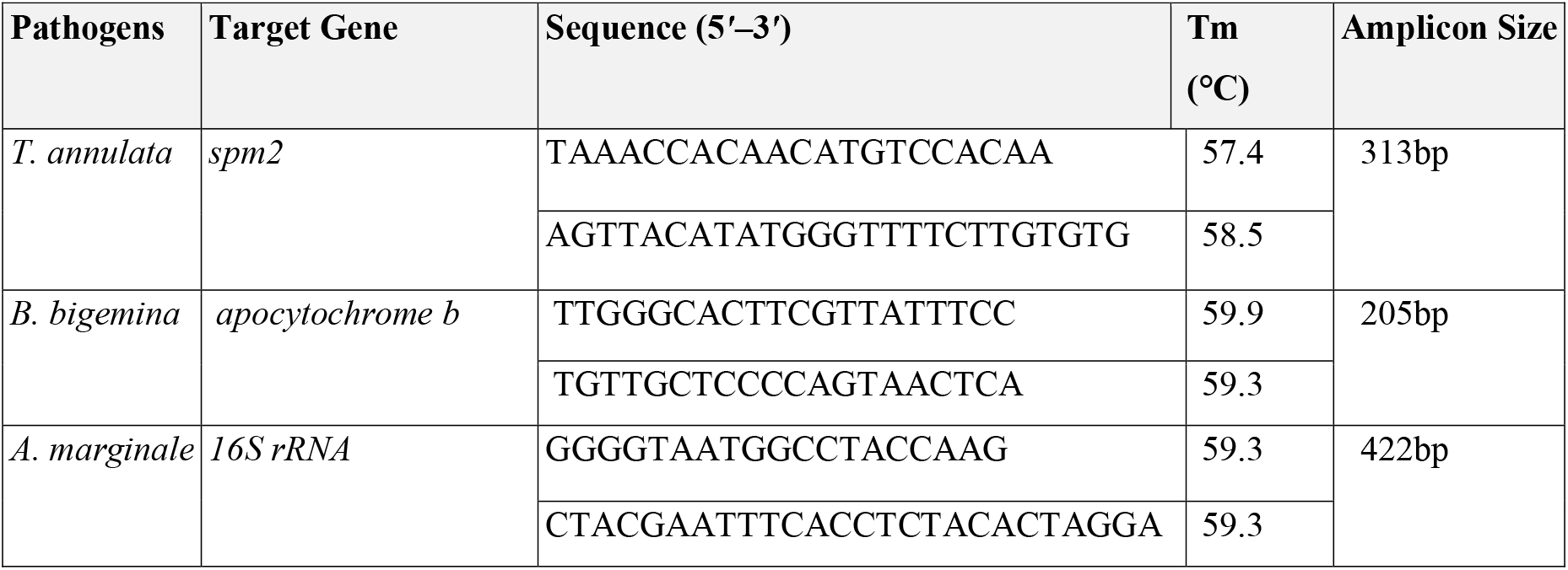
List of primers designed from target genes for simultaneous diagnosis of *T. annulata, A. marginale* and *B. bigemina* by multiplex PCR

### Multiplex PCR

The thermal PCR program was standardized for multiplex PCR to detect the genomic DNA of three haemo-parasite (*T. annulata, B. bigemina, and A. marginale*) in a single run using DNA extracted from cattle blood samples. All isolated DNA samples from cattle blood that was found infected with single or multiple haemo-parasite by GSBS was used to run the multiplex PCR.

Briefly, PCR assay was performed in a total volume of 25 μl containing 3 μl (50 ng/μl) of the genomic DNA and 2 μl primers each of forward and reverse primer prepared after mixing of equal volume of 10 micro-molar working solution of the three forward and reverse primers and 12.5 μl of OneTaq Hot Start master mix with buffer (Imperial Life Sciences (P) Limited), and ultra-pure nuclease free water to make the final volume. Amplification involved a hot start of 5 min at 95°C, followed by 35 cycles of 1 min at 95°C, 1 min at 58°C, 1 min at 72°C, and a final extension step of 10 min at 72°C.

The amplified PCR products were subjected to electrophoresis on 2% agarose gel stained with ethidium bromide, visualized under UV light, and photographed in trans-illuminator apparatus. The amplicons molecular sizes were estimated by including a Quick-Load 100 bp DNA Ladder (Imperial Life Sciences Private Limited) and visualised and documented on Gel-Doc (Azure Biosystems c200). DNA positive controls of *T. annulata* were kindly provided by NAIB, Hyderabad, while positive control of *A. marginale* (MK834271) and *B. bigemina* (MH936010) was used from DNA found positive after specific amplification and sequencing for species specific amplified products. Distilled water served as negative control.

### Statistics

Data were analysed for mean and standard error. Analysis of variance (ANOVA) using post hoc LSD test of significance, if any, was observed at P<0.05. The observation of GSBS and Multiplex PCR was compared for sensitivity (S_e_), specificity (S_p_), positive predictive values (PPV) and negative predictive value (NPV) by the method elaborated earlier (Trevethan 2017). Cohen’s kappa coefficient was calculated by the method described (Landis and Koch 1977).

## Results

The study indicated that the endemic region had high (76.67%) infection of tick transmitted haemo-parasite in selected crossbred cattle based on conventional Giemsa blood smear examination (Fig. 1). The infected animals had clinical signs such as high fever, drop in milk yield, inappetance to anorexia, high percentage (30.44) of infected cattle were co-infected with more than one haemo-parasite. The haematological changes observed in these cattle are indicative of significant RBCs destruction, anaemia (reduced Hb level) and leukopenia in infected cattle compared to non-infected cattle (Table 2). The study corroborated with earlier reports of these changes associated with tick transmitted haemo-parasite in cattle (Ganguly et al. 2017; Larcombe et al. 2019; Prabhakaran et al. 2021) and buffaloes (Kumar et al. 2021). Molecular confirmation based on PCR amplification requires multiple attempts to detect co-infected samples. Multiplex PCR based on species specific primers was observed to be useful and 100 % specific in detecting the co-infection of tick transmitted haemo-parasite in a single run (Table 3). The visualization was clear and specific on gel doc examination (Fig. 2). Negative samples did not result in any amplification. Further, the results of multiplex PCR considered as confirmatory was compared with results of GSBS. The unmatched results of GSBS was considered as false positive and negative. The sensitivity (S_e_), specificity (S_p_), positive predictive values (PPV), negative predictive values (NPV) and Cohen’s kappa coefficient of the GSBS observation with multiplex PCR are depicted in table 4. The S_p_, PPV and kappa coefficient of agreement was highest for diagnosis of *B. bigemina* and lowest for *A. marginale*. The S_e_ was highest for detection of *T. annulata* and lowest for *B. bigemina*. The overall Kappa coefficient for diagnosis of these parasites by the both the test are in moderate (0.56) agreement, though has high S_e_ (78.9), S_p_ (76.5), and PPV (78.9) percentage. This the multiplex PCR is able to detect false negative samples based on GSBS due to low level of parasitaemia and also able to rectify false positive results due to differences in subjective observations.

**Figure 1.**
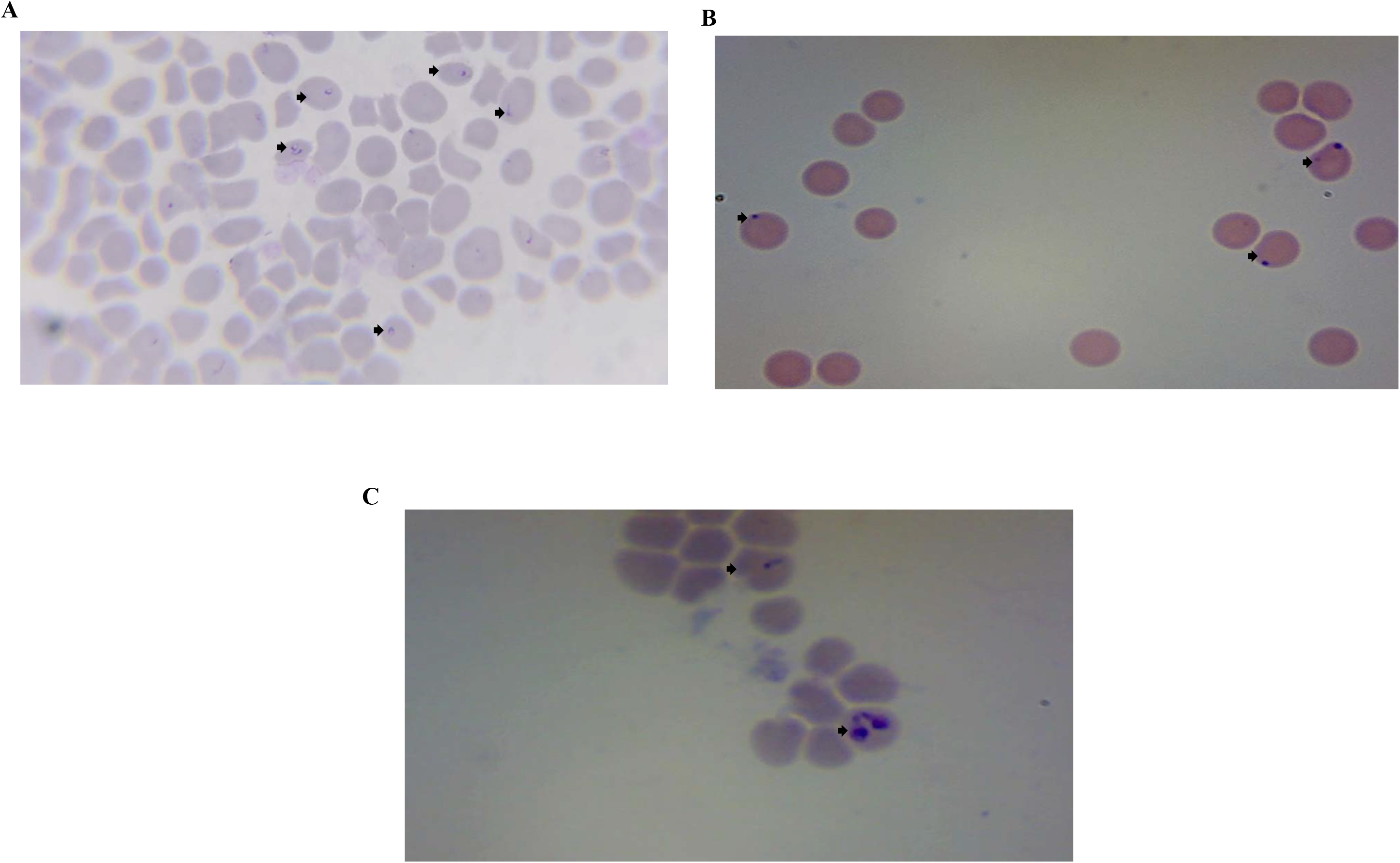
Diagnosis based on conventional GSBS. Representative photographs were taken under oil immersion microscope 1000x magnification. **A**. Infected RBCs (Arrow) with *T. annulata* prioplasm; **B**. Infected RBCs with A. marginale (arrow); C. Infected RBCs with *B. bigemina* (paired pear-shaped bodies) and *T. annulata* (above arrow)

**Table 2.** Haematology of crossbred cattle diagnosed for tick transmitted pathogens based on GSBS

| GSBS findings | Haemoglobin (g/L) | RBC count ($\times 10^6/\mu\text{L}$) | WBC count ($\times 10^3/\mu\text{L}$) | PCV (%) |
| --- | --- | --- | --- | --- |
| *T. annulata* (n=7) | 7.39<sup>a</sup>±0.43<br>(3.50-5.70) | 4.20<sup>a</sup>±0.20<br>(1.64-3.24) | 6.45<sup>a</sup>±0.45<br>(5.12-8.21) | 14.71<sup>a</sup>±1.46<br>(9.0-21.0) |
| *A. marginale* (n=9) | 7.69<sup>a</sup>±0.44<br>(5.8-9.6) | 5.54<sup>b</sup>±0.28<br>(3.98-6.42) | 5.99<sup>a</sup>±0.31<br>(4.89-7.12) | 15.67<sup>a</sup>±1.50<br>(9.0-22.0) |
| Co-infections (n=7) | 6.74<sup>a</sup>±0.37<br>(5.8-8.3) | 4.54<sup>a</sup>±0.23<br>(3.88-5.62) | 5.57<sup>a</sup>±0.31<br>(4.66-6.84) | 12.43<sup>a</sup>±1.29<br>(9.00-18.0) |
| Negative (n=7) | 10.94<sup>b</sup>±0.23<br>(9.9-11.8) | 7.78<sup>c</sup>±0.34<br>(6.24-8.92) | 7.85<sup>b</sup>±0.41<br>(6.38-9.45) | 26.29<sup>b</sup>±0.75<br>(23.0-29.0) |
| Reference Range (Jain, 1993) | **80-150** | **5-10** | **4-12** | **24-46** |

Values in column with different superscript varies significantly (P<0.05)

**Table 3.** Comparison of observation based on conventional GSBS examination and multiplex PCR developed

| Sample No | GSBS |  |  | Multiplex PCR |  |  |
| --- | --- | --- | --- | --- | --- | --- |
|  | A | T | B | A | T | B |
| 1 | +ve | -ve | -ve | +ve | +ve | -ve |
| 2 | +ve | +ve | -ve | +ve | +ve | -ve |
| 3 | +ve | +ve | -ve | +ve | +ve | -ve |
| 4 | +ve | +ve | -ve | +ve | +ve | -ve |
| 5 | -ve | +ve | -ve | +ve | +ve | -ve |
| 6 | +ve | -ve | -ve | -ve | -ve | +ve |
| 7 | -ve | +ve | +ve | -ve | +ve | +ve |
| 8 | -ve | +ve | +ve | -ve | +ve | +ve |
| 9 | +ve | -ve | +ve | -ve | -ve | +ve |
| 10 | -ve | +ve | -ve | -ve | -ve | +ve |
| 11 | +ve | +ve | -ve | -ve | +ve | -ve |
| 12 | +ve | -ve | -ve | +ve | -ve | -ve |

Where, A= *A. marginale*; B= *B. bigemina*; T= *T. annulata*; +ve: Positive; -ve: Negative

**Figure 2.**
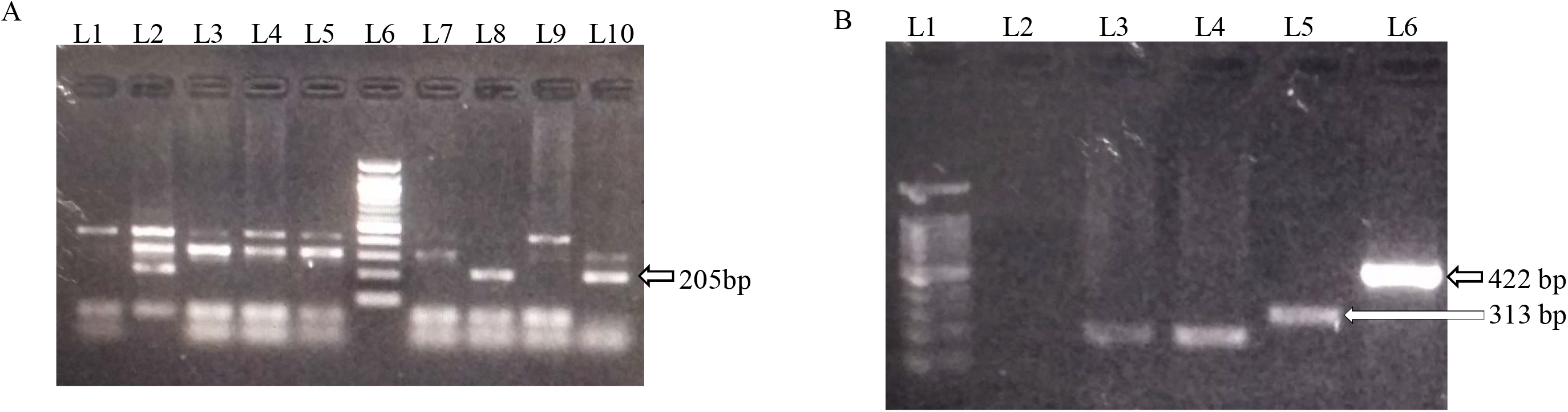
Multiplex PCR product agarose gel electrophoresis visualized under UV light of Gel Doc with positive amplification at 205, 313 and 422 bp for *B. bigemina, T. annulata* and *A. marginale*, respectively. Lane 1: Positive control; Lane 5: 100 bp Ladder; Remaining lanes: Samples 1-8; **B**. Lane 1:100 bp ladder; Lane 2: Negative control; Remaining lanes: Samples 9-12

**Table 4.** Comparison of sensitivity (S_e_), specificity (S_p_), positive predictive values (PPV), negative predictive values (NPV) and Cohen’s kappa coefficient of the GSBS observation with multiplex PCR

| Pathogens | $S_e$ (%) | $S_p$ (%) | PPV (%) | NPV (%) | Kappa Coefficient |
| --- | --- | --- | --- | --- | --- |
| A | 83 | 50 | 62.5 | 75 | 0.34 |
| T | 87.5 | 75 | 87.5 | 75 | 0.62 |
| B | 60 | 100 | 100 | 22.2 | 0.63 |
| A+T+B | 78.9 | 76.5 | 78.9 | 76.5 | 0.56 |

Where, A= *A. marginale*; B= *B. bigemina*; T= *T. annulata*; Kappa coefficient of 0.6 and above is considered as in agreement

## Discussion

Tick transmitted diseases with high infection in bovine population of Bihar with *Theileria* spp., *A marginale*, and *B. bigemina* and their co-infection has been frequently reported (Kala et al. 2018; Kumar et al. 2019; Kumar et al. 2021; Prabhakaran et al. 2021; Roy 2021). Our finding also suggest high rate of infection of these hemo-parasite and association of more than one type parasite in the animals screened by GSBS. These results have few limitation and dependent on expert examination and staining procedures (Prabhakaran et al. 2021). It was observed that cattle infected with these parasites adversely affected haematological parameters. These changes are associated with capability of infecting parasite to reach the circulation and invade the RBCs and corroborates with earlier findings from mixed infected cattle of southern India (Jayalakshmi et al. 2019). Severity of changes varied depending upon the type and degree of RBCs infected by these parasites. The confirmatory molecular diagnostic method was developed to detect co-infected state in animal in single run by multiplex PCR using specific gene primers which was found useful and corroborates with the findings of GSBS. *Spm2 gene* used for detection of *T. annulata* has not been attempted earlier for multiplex PCR development and has been reported to be expressed both in sporozoites and in later stages (macroschizont-infected leucocytes and piroplasms) of life-cycle of *T. annulata* (Knight et al. 1998). *Apocytochrome b* gene has been reported to be very effective for detection of *B. bigemina* and *B. bovis* by targeting a species-specific region of this gene (Ganzinelli et al. 2020). Similar methods of multiplex PCR are also reported based on different target genes. Kundave et al., (2018) developed multiplex PCR using *Tams1, 18S rRNA* and *16S rRNA* genes of *T. annulata, B. bigemina* and *A. marginale*, respectively. Bilgiç et al., (2013) developed a multiplex PCR assay for simultaneous detection of *T. annulata, A. marginale* and *B. bovis* in cattle using *cytochrome b* gene, *major surface protein–1β* encoding gene, and *VESA–1α* gene, respectively. The results of S_e_, S_p_, PPV, NPV and Cohen’s kappa coefficient of the GSBS observation with multiplex PCR indicates that diagnosis of *A. marginale* is difficult with GSBS attributed to similarity of *A. marginale* with staining artifacts and sometimes polymorphic attributes of *T. annulata*. On the contrary, diagnosis of *B. bigemina* had high S_p_, PPV which is attributed to the sampling of animals mostly after characteristic clinical symptoms of red urine. The Landis and Koch (1977) have proposed the following as standards for strength of agreement for the kappa coefficient: ≤0=poor, 0.01–0.20=slight, 0.21–0.40=fair, 0.41–0.60=moderate, 0.61–0.80=substantial, and 0.81-1=almost perfect. The Cohen’s kappa coefficients indicates that diagnosis of *T. annulata* and *B. bigemina* are in substantial agreement with the GSBS observation with obvious reason of sampling from animal with characteristic though overlapping clinical symptoms.

## Conclusion

A new method of multiplex PCR has been developed for simultaneous detection of *T. annulata, B. bigemina* and *A. marginale* in cattle in a single run. This methods will be useful for researchers for diagnosis of co-infection of tick transmitted diseases in cattle in endemic areas where possibilities of co-infection exists. Further refinement in the methodology needs to be undertaken to diagnose other parasites like *T. orientalis, B. bovis, A. centrale*, etc.

## Author Contributions

PK, MK conceived the study, designed the primers, KS supervised the study PK, RRK collected samples from field and initial screening. PK, PS and MK standardized the technique of multiplex PCR. AK, PK, RRK and PS analysed the data statistically. PK, RRK, MK, PS and AK wrote the manuscript.

## Acknowledgements

Authors acknowledge support and cooperation of Director, ICAR-Research Complex for Eastern Region, Patna for providing necessary funds and facilities for undertaking this intra-mural project. Support provided by project staff of ICAR Outreach program on Zoonotic Diseases is also duly acknowledged.

